# Ebola virus glycoprotein domains associated with protective efficacy

**DOI:** 10.1101/2021.05.22.445257

**Authors:** Bharti Bhatia, Wakako Furuyama, Thomas Hoenen, Heinz Feldmann, Andrea Marzi

## Abstract

Ebola virus (EBOV) is the cause of sporadic outbreaks of human hemorrhagic disease in Africa, and the best-characterized virus in the filovirus family. The West Africa epidemic accelerated the clinical development of vaccines and therapeutics leading to licensure of vaccines and antibody-based therapeutics for human use in recent years. The most widely used vaccine is based on vesicular stomatitis virus (VSV) expressing the EBOV glycoprotein (GP)(VSV-EBOV). Due to its favorable immune cell targeting, this vaccine has also been used as base-vector for the development of second generation VSV-based vaccines against Influenza, Nipah, and Zika viruses. However, in these situations it may be beneficial if the immunogenicity against EBOV GP is minimized to induce a better protective immune response against the other foreign immunogen. Here, we analyzed if EBOV GP can be truncated to be less immunogenic yet still able to drive replication of the vaccine vector. We found that the EBOV GP glycan cap and the mucin-like domain are both dispensable for VSV-EBOV replication. The glycan cap domain, however, appears critical for mediating a protective immune response against lethal EBOV challenge in mice.

## Introduction

Filoviruses were identified in 1967 and have become known to cause severe human disease with case fatality rates of up to 90% [1]. The family *Filoviridae* is divided into six genera and 11 species and includes several human-pathogenic viruses [2]. Ebola virus (EBOV) and Marburg virus are the best-known members of this virus family because infrequent spill-over events into the human population with subsequent human-to-human transmission cause outbreaks of Marburg virus and Ebola virus disease (EVD) [3]. EBOV was headlining the news when the West African countries Guinea, Sierra Leone and Liberia were facing an EVD epidemic with over 28,000 cases with over 11,000 fatalities [4]. During this epidemic the clinical development of vaccine and therapeutic candidates was accelerated resulting in the approval of an EBOV vaccine by the United States Food and Drug Administration (FDA) and by the European Medicines Authority (EMA) in 2019 [5,6]. This licensed, live-attenuated vaccine is based on vesicular stomatitis virus (VSV); its glycoprotein was replaced with the EBOV glycoprotein GP, which is the main immunogen of the virus [7]. The vaccine VSV-EBOV, also known as rVSV-ZEBOV and marketed under the brand name Ervebo^®^, has been shown to protect nonhuman primates (NHPs) from lethal disease after administration of a single dose [8]. Mechanistic studies revealed that antibodies specific to the EBOV GP are the main mediators of protection [9], however, the fast-acting nature of the vaccine is likely due to a combination of strong innate followed by adaptive immune responses [10].

In recent years, we have developed second generation vectors based on VSV-EBOV. The concept is founded on the favorable immune cell targeting of the EBOV GP which has been hypothesized to be important for the fast-acting nature of VSV-EBOV [10,11]. VSV-EBOV-based vectors have been successfully developed as vaccine candidates for a number of different viruses including influenza, Nipah (NiV) and Zika viruses (ZIKV) [12–14]. Most recently, a vaccine against SARS-CoV-2 was developed based on this vector which quickly protected NHPs from COVID-19 [15]. These vaccines express an additional viral immunogen like the ZIKV pre-matrix and envelope proteins, and induce protective responses against challenge with both EBOV and ZIKV [13]. However, the strong immunogenicity of the EBOV GP may negatively impact the immune responses directed to the second immunogen. Alternatively, second generation VSV-EBOV-based vectors can be produced as true bivalent vaccine vectors inducing similar protective efficacy in parallel against challenge with EBOV and another pathogen.

Here, we investigate if the immunogenicity of the EBOV GP can be reduced without compromising vector replication. For this we generated VSV-EBOV vectors expressing GPs harboring deletions of the two most immunogenic domains, the mucin-like domain (MLD) and the glycan cap (GC) (Fig. 1A). We found that all vectors replicated well *in vitro*, however, protective efficacy against lethal challenge in the EBOV mouse model was reduced when the GC was lacking. Our data suggests that the GC of the EBOV GP is associated with protective immunity of GP-based EBOV vaccines.

**Figure 1.**
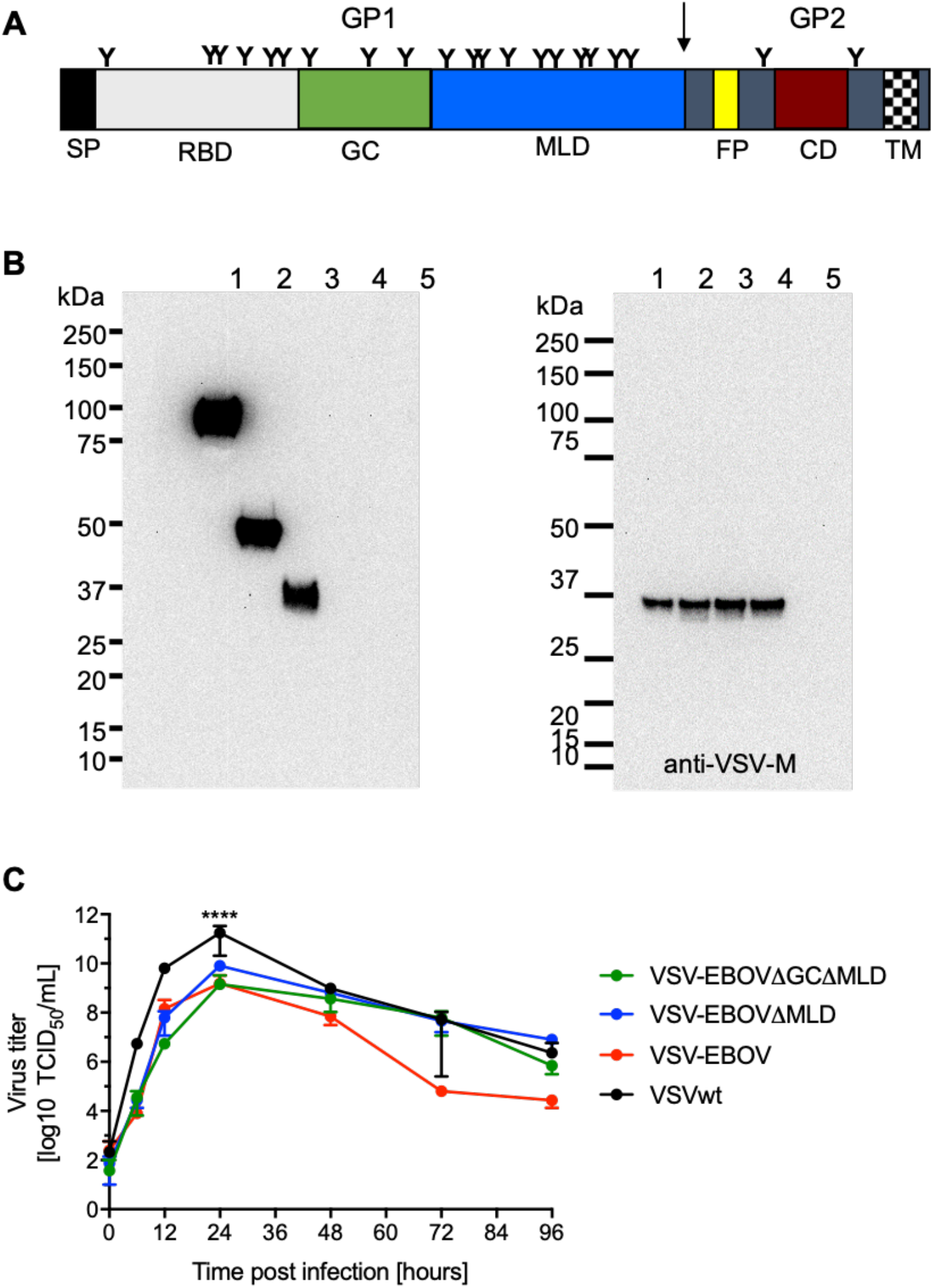
Vaccine construction and *in vitro* characterization. **(A)** Schematic of the EBOV glycoprotein GP. SP signal peptide; RBD receptor-binding domain; GC glycan cap; MLD mucin-like domain; FP fusion peptide; CD coiled-coil domain; TM transmembrane domain. Arrow indicates furin-cleavage site. **(B)** The VSV-EBOV vector was modified to express a GP containing a deletion of the MLD (VSV-EBOVΔMLD) or the GC plus MLD (VSV-EBOVΔGCΔMLD). After successful recovery of the viruses from plasmid transfections, antigen expression was confirmed by Western blot analysis using monoclonal antibodies specific for the EBOV GP (left panel) or VSV matrix (M) protein (right panel). Lane 1 VSV-EBOV; lane 2 VSV-EBOVΔMLD; lane 3 VSV-EBOVΔGCΔMLD; lane 4 VSV wildtype (wt); lane 5 uninfected control. **(C)** Growth kinetics were performed in triplicate on Vero E6 cells at a multiplicity of infection of 0.01. Geometric mean and SD are depicted. Statistical significance is indicated.

## Materials and Methods

### Ethics statement

All infectious work was performed in the maximum containment laboratory (MCL) at the Integrated Research Facility, Rocky Mountain Laboratories (RML), Division of Intramural Research (DIR), National Institute of Allergy and Infectious Disease (NIAID), National Institutes of Health (NIH). Vaccinations were carried out in the biosafety level 2; EBOV was handled exclusively in the MCL according to standard operating procedures (SOPs) approved by the Institutional Biosafety Committee (IBC). The animal work was approved by the Institutional Animal Care and Use Committee (IACUC) and performed according to the guidelines of the Association for Assessment and Accreditation of Laboratory Animal Care, International and the Office of Laboratory Animal Welfare. All procedures on animals were carried out by trained personnel following SOPs approved by the IBC. Humane endpoint criteria in compliance with IACUC-approved scoring parameters were used to determine when animals should be humanely euthanized.

### Cells and viruses

African green monkey kidney (Vero E6) cells were grown in DMEM (Sigma-Aldrich) containing 2% or 10% fetal bovine serum (FBS), 2 mM L-glutamine, 50 U/mL penicillin, and 50 μg/mL streptomycin (all from Thermo Fisher Scientific). Baby hamster kidney (BHK) - T7 cells were grown in DMEM containing 10% tryptose phosphate broth (Thermo Fisher Scientific), 5% FBS, L-glutamine, penicillin, and streptomycin. THP-1 cells were grown in RPMI (Sigma-Aldrich) containing 10% fetal bovine serum (FBS), L-glutamine, penicillin, and streptomycin. VSV-EBOV lacking the MLD (VSV-EBOVΔMLD; deletion of aa 309-489 [16]) was previously described [17]. A VSV vector expressing EBOV GP lacking the GC and MLD VSV-EBOVΔGCΔMLD (deletion of aa 228-489) was constructed and recovered from plasmid as previously described [13]. VSV wildtype (VSVwt) and VSV-EBOV were used as control vaccines [18]. Mouse-adapted EBOV (MA-EBOV) was used for the challenge study in mice [19]. All viruses were propagated and titered on Vero E6 cells, sequenced confirmed, and stored at −80 °C.

### Western blot analysis

Vero E6 cells were either mock-infected or infected with the different recombinant VSV vectors (MOI 0.1). Cell culture supernatant was mixed 1:1 (v/v) with sodium dodecyl sulfate-polyacrylamide (SDS) gel electrophoresis sample buffer containing 20% β-mercaptoethanol and heated to 99 °C for 10 min. Proteins were separated on SDS-PAGE using TGX criterion pre-cast gels (Bio-Rad Laboratories). Subsequently, proteins were transferred to a Trans-Blot polyvinylidene difluoride membrane (Bio-Rad Laboratories). The membrane was blocked for 3 h at room temperature in PBS with 3% powdered milk and 0.05% Tween 20 (Thermo Fisher Scientific). Protein detection was performed using the anti-EBOV GP (ZGP 12/1.1, 1μg/ml; kindly provided by Ayato Takada, Hokkaido University, Sapporo, Japan), and anti-VSV matrix (VSV-M) (23H12, 1:1,000; Kerafast Inc.) antibodies. After staining with the anti-mouse horse-radish peroxidase (HRP)-labeled secondary antibody at 1:10,000 (cat. #715-035-151, Jackson ImmunoResearch), the blots were imaged using the SuperSignal West Pico chemiluminescent substrate (Thermo Fisher Scientific) and a FluorChem E system (Protein simple).

### Growth kinetics

Vero E6 cells were grown to confluency in a 12-well plate and infected in triplicate with VSVwt, VSV-EBOV, VSV-EBOVΔMLD, and VSV-EBOVΔGCΔMLD (MOI of 0.01). The inoculum was removed, cells were washed 3 times with DMEM, and covered with DMEM containing 2% FBS. Supernatant samples were collected at 0, 6, 12, 24, 48, 72, and 96 hours post-infection and stored at −80 °C. The titer of the supernatant samples was determined performing a median tissue culture infectious dose (TCID_50_) assay on Vero E6 cells as previously described [20].

### Vaccination and protective efficacy in mice

Female CD1 mice (5-6 weeks old) were obtained from Envigo (Somerset, NJ). Groups of 14 mice were vaccinated intraperitoneally (IP) on day -28 with 1 × 10^4^ PFU (2 sites, 0.1 ml each) with VSVwt, VSV-EBOV, VSV-EBOVΔMLD, or VSV-EBOVΔGCΔMLD. Unvaccinated/naïve animals served as a control group. On the day of challenge (day 0) 4 animals from each group were euthanized for blood collection. The remaining 10 animals in each group were challenged IP with 1,000 median lethal dose (LD_50_)(10 focus-forming units) of MA-EBOV [21]. On day 4 after challenge, 4 mice in each group were euthanized for blood and tissue sample collection. Samples were stored at −80 °C until titration as previously described [22]. Surviving mice were monitored until 42 days post-infection when a single, terminal blood sample was collected.

### Enzyme-linked immunosorbent assay (ELISA)

Serum samples from MA-EBOV-infected mice were inactivated by γ-irradiation (4 megarads) [23] and used in BSL2 according to IBC-approved SOPs. For the detection of EBOV GP-specific antibodies in mouse sera, soluble EBOV GP lacking the transmembrane domain (EBOV GPΔTM) was produced and used in an enzyme-linked immunosorbent assay (ELISA) as described before [24]. The reaction was measured using the Synergy™ HTX Multi-Mode Microplate Reader (BioTek).

### Neutralization assay

Transcription and replication competent virus-like particles (trVLPs) were produced in HEK293 cells as described previously [25]. In place of the T7-driven tetracistronic minigenome plasmid an RNA polymerase II-driven tetracistronic EBOV minigenome plasmid (pCAGGS-4cis-EBOV-eGFP) expressing GFP as reporter and hammerhead and hepatitis delta virus ribozymes flanking the minigenome was used (details about the cloning strategy will be provided upon request). The mouse sera were heat-inactivated for 30 min at 50°C before use. For neutralization assays, trVLPs were appropriately diluted to 100-300 infectious units and then incubated for 60 min at room temperature with serial dilutions of the mouse serum samples. After incubation, the mixture was inoculated onto Vero E6 cells. After 72 hours, neutralizing activity was quantified by counting the number of GFP-positive cells. The relative percentage of infectivity was calculated by setting the number of cells inoculated with trVLPs in the presence of negative control mouse serum to 100%.

### Statistical analysis

Statistical analysis was performed using Prism 8 (GraphPad). Survival curves were analyzed using Mantel-Cox test. Mouse body weight changes were analyzed using 2-way ANOVA with Tukey’s multiple comparison. Viral and ELISA titers were analyzed with the Mann-Whitney test. Statistically significant differences are indicated as follows: p<0.0001 (****), p<0.001 (***), p<0.01 (**), and p<0.05 (*).

## Results

### VSV vectors expressing EBOV GP with deletions replicate well in cell culture

The VSV-EBOVΔGCΔMLD vector was generated by deleting aa 227-489 encoding the GC and MLD (ΔGCΔMLD) (Fig. 1A) in the EBOV GP and cloning this open reading frame into the VSV vector as previously described [18]. After recovery of the virus from plasmid following an established protocol [13], the sequence was confirmed. The presence of the EBOV GPΔGCΔMLD on VSV particles was analyzed by Western blot analysis from infected VeroE6 cell culture supernatants in comparison to previously rescued VSVwt, VSV-EBOV [18] and VSV-EBOVΔMLD [17] (Fig. 1B). An antibody specific to the EBOV GP confirmed expression and incorporation of similar levels of EBOV GP, EBOV GPΔMLD and EBOV GPΔGCΔ into recombinant VSV particles (Fig. 1B, left panel). Detection of similar levels of VSV-M in all preparations indicated the use of similar amounts of recombinant VSV particles (Fig. 1B, right panel). Next, we examined the growth kinetics of the recombinant VSVs expressing the different versions of EBOV GP. We found no significant differences in virus growth or endpoint titers between VSV-EBOV, VSV-EBOVΔMLD and VSV-EBOVΔGCΔMLD (Fig. 1C). VSVwt had a significantly higher titer 24 hours after infection indicating *in vitro* attenuation of the recombinant VSVs expressing different forms of the EBOV GP.

### VSV-EBOVΔGCΔMLD failed to protect mice from lethal EBOV challenge

The protective efficacies of VSV vaccines expressing different version of EBOV GP were evaluated in the well-established mouse model for EBOV using MA-EBOV infection of CD1 mice [13,21]. Mice were IP vaccinated with a single dose of 1 × 10^4^ PFU 28 days before challenge (Fig. 2A). On the day of challenge (day 0), 4 mice in each group were euthanized for serum collection and the remaining mice were all IP infected with 1,000 LD_50_ of MA-EBOV (Fig. 2A). Mice were monitored daily, and body weights were collected for the first 14 days (Fig. 2B). Starting on day 3 post-challenge, mice vaccinated with VSV-EBOVΔGCΔMLD or VSVwt and unvaccinated controls showed initial signs of disease including ruffled fur and weight loss (Fig. 2B). Over the next 5 days all mice in the VSVwt and control groups succumbed to the infection, as did 5/6 mice in the VSV-EBOVΔGCΔMLD group (Fig. 2C). Mice vaccinated with VSV-EBOV or VSV-EBOVΔMLD never showed signs of disease and survived indicating complete protection (Fig. 2B, C). On day 4 post-challenge, 4 mice per group were euthanized for sample collection to determine virus loads and antibody levels (Fig. 2A). We found that the virus replicated to high titers in the blood, liver and spleen of mice vaccinated with VSV-EBOVΔMLD or VSVwt as well as unvaccinated controls (Fig. 2D). In contrast, no virus was isolated from mice vaccinated with VSV-EBOV with the exception of one single spleen sample (Fig. 2D). Mice vaccinated with VSV-EBOVΔMLD had replicating virus in blood, liver and spleen samples albeit to a significantly lower titer compared to animals in the VSV-EBOVΔMLD, VSVwt or unvaccinated control groups that succumbed to MA-EBOV challenge (Fig. 2D). Surviving mice were euthanized on day 42 (Fig. 2A).

**Figure 2.**
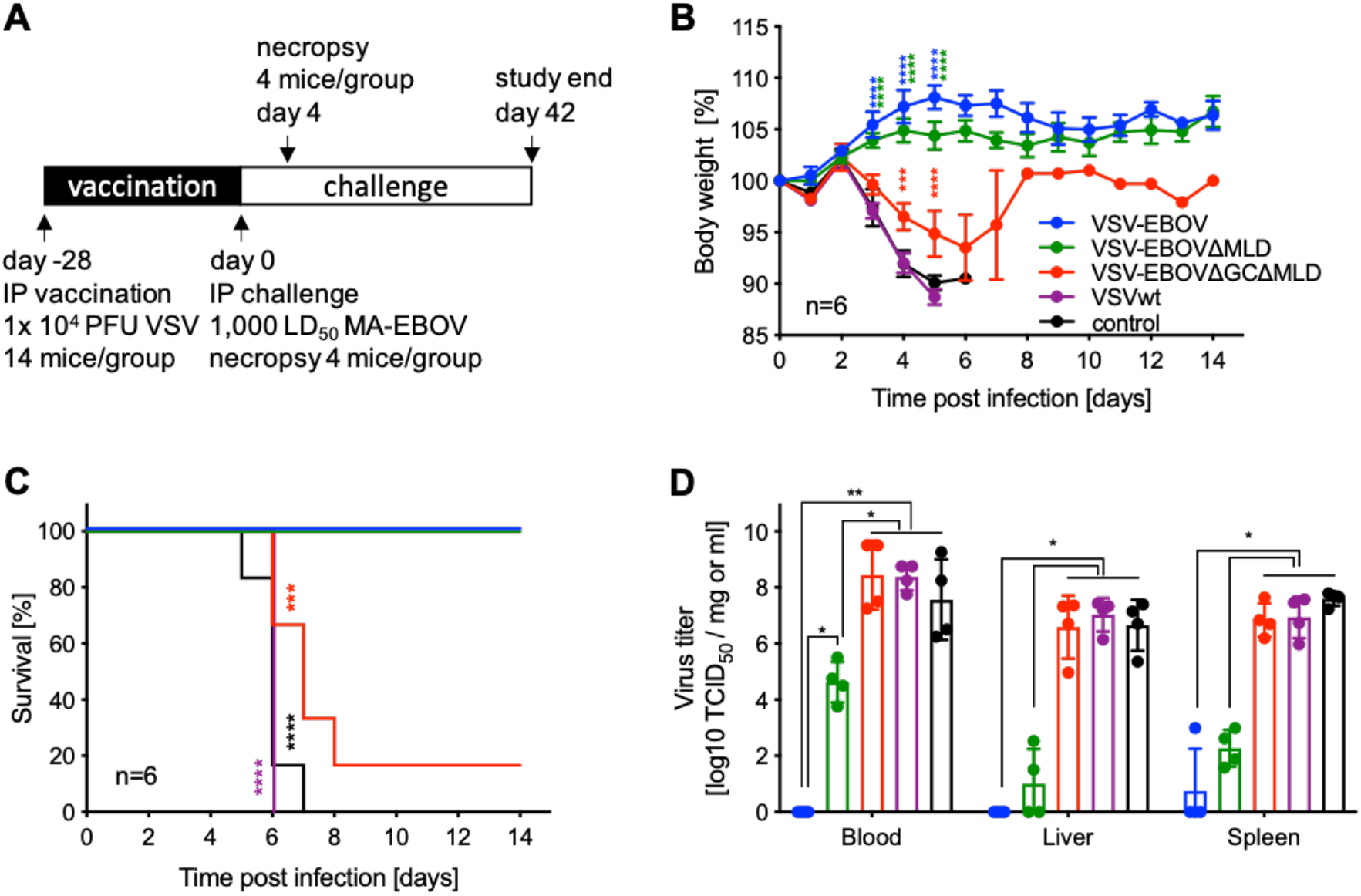
Study outline and protective efficacy in mice. **(A)** Study outline. Groups of CD1 mice were intraperitoneally (IP) vaccinated with a single dose of 1 × 10^4^ PFU of the vaccines 28 days prior to lethal challenge. **(B)** Body weight changes and **(C)** survival curves are shown. **(D)** On day 4 after challenge, 4 mice in each group were euthanized for sample collection, and virus titers were determined. Geometric mean and SD are depicted. Statistical significance is indicated.

### Serum antibody levels after vaccination predict survival

The amounts of EBOV GP-specific IgG were determined in serum samples on day 0 (28 days after vaccination; n=4), day 4 post-challenge (n=4) and all surviving mice (VSV-EBOV, VSV-EBOVΔMLD n=6; VSV-EBOVΔGCΔMLD n=1). On day 0, mice vaccinated with any of the VSVs expressing a version of EBOV GP had significant EBOV GP-specific titers compared to VSVwt-vaccinated or unvaccinated control mice (Fig. 3A). The EBOV GP-specific titers in sera of mice vaccinated with VSV-EBOVΔGCΔMLD were lower on day 0 compared to the VSV-EBOV or VSV-EBOVΔMLD groups, however, the difference was not statistically significant (Fig. 3A). On day 4, the level of EBOV GP-specific IgG in the serum of VSV-EBOVΔGCΔMLD-vaccinated mice was comparable to control groups and significantly lower to the other EBOV GP-based vaccines (Fig. 3A).

**Figure 3.**
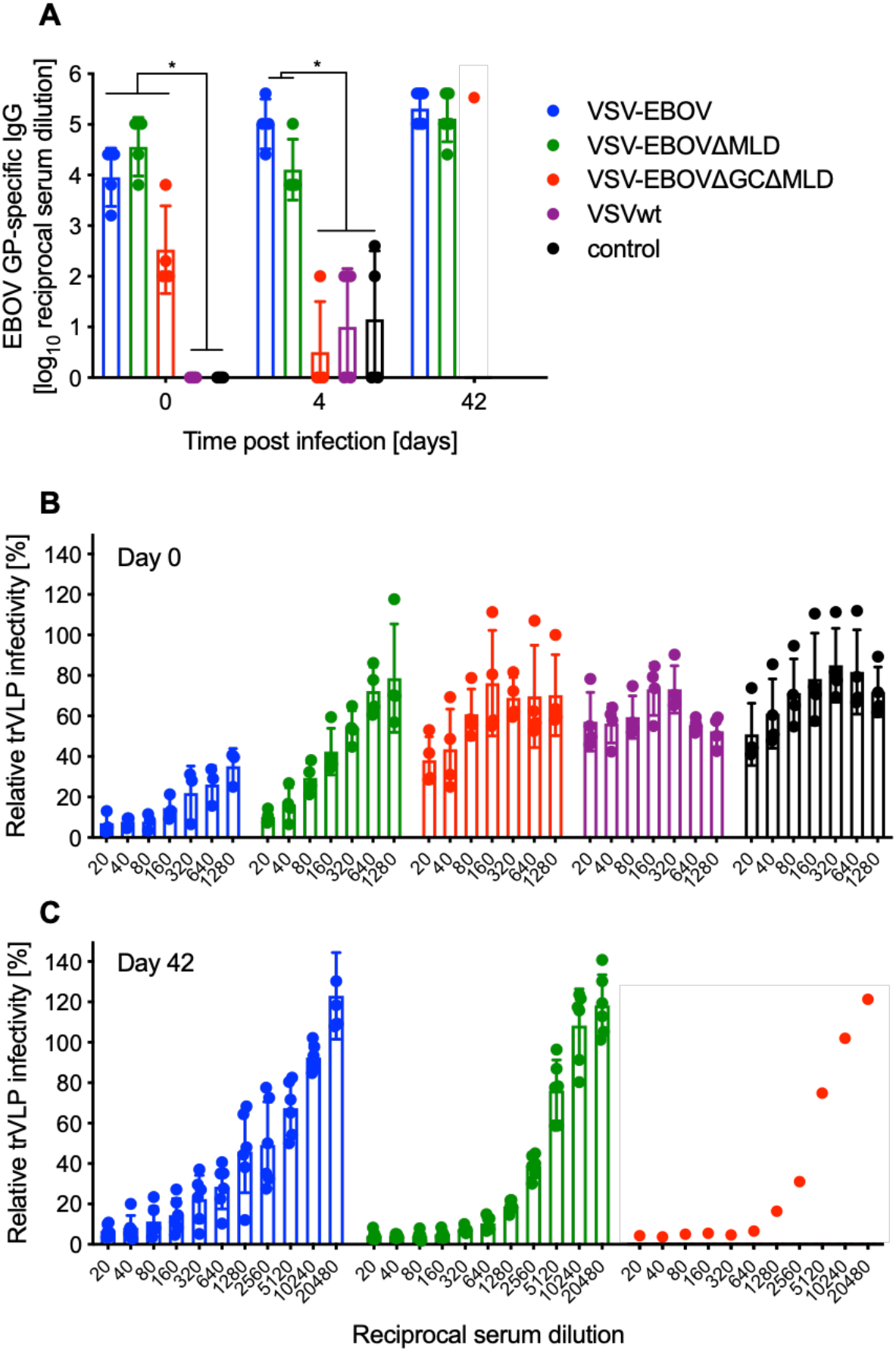
Humoral immune responses in vaccinated and challenged mice. **(A)** EBOV GP-specific IgG was assessed in serum samples from mice on day 0 (n=4; 28 days after vaccination), day 4 (n=4) and day 42 (all survivors). Geometric mean and SD are depicted. Statistical significance is indicated. **(B)** Neutralizing activity in the serum of vaccinated mice (day 0, n=4 per group) and challenge survivors (day 42; n=6 for VSV-EBOV and VSV-EBOVΔMLD or n=1 for VSV-EBOVΔGCΔMLD) **(C)** was determined using the EBOV trVLP system expressing eGFP. Mean and SD are depicted.

Next, we determined differences in neutralizing antibody titers using a well-characterized surrogate system [26]. Similar to the EBOV GP-specific IgG ELISA data, the neutralization is strongest after vaccination with VSV-EBOV, followed by VSV-EBOVΔMLD (Fig. 3B). Mice vaccinated with VSV-EBOVΔGCΔMLD, VSVwt and unvaccinated controls had limited to no neutralizing activity in their serum (Fig. 3B). Neutralization at 50% occurred at the following serum concentrations VSV-EBOV >1:1280, VSV-EBOVΔMLD ~1:160, VSV-EBOVΔGCΔMLD ~1:20.

Sera of VSVwt and unvaccinated control animals were non-neutralizing (Fig. 3B). We also examined the neutralizing activity in serum samples collected from the surviving mice and found that the neutralizing titers for the VSV-EBOV group were comparable to day 0 (Fig. 3C). However, the neutralizing activity in surviving mice vaccinated with VSV-EBOVΔMLD and the single surviving animal vaccinated with VSV-EBOVΔGCΔMLD was clearly boosted compared to day 0 indicating MA-EBOV replication following challenge (Fig. 3C). The boosting effect through challenge was less clear with animals vaccinated with VSV-EBOV indicating more potent protection (Fig. 3C).

## Discussion

VSV-EBOV was approved by the US FDA and the EMA for human use in 2019 and as of February 2020 has been administered to over 290,000 people [27]. This number is growing with the latest EBOV outbreaks occurring in the Democratic Republic of the Congo and Guinea (https://www.cdc.gov/vhf/ebola/outbreaks/index-2018.html) [28]. Because of its ease of construction, production and efficacy in preventing disease, VSV-EBOV has been pursued as a base vector for the development of vaccines against other emerging viruses including influenza, NiV and ZIKV [12–14]. In fact, the clinical development of a VSV-NiV vaccine based on VSV-EBOV expressing NiV G has been funded by the Coalition for Epidemic Preparedness Innovations(CEPI) and is well on its way [29].

It was previously shown that pre-existing immunity to the VSVΔG vector does not hinder re-vaccination with another VSVΔG-based vaccine [30]. Little is known, however, about the influence of pre-existing immunity to EBOV or EBOV GP on vaccine efficacy using VSV-EBOV or VSV-EBOV-based vaccine candidates. Vaccination utilizing EBOV GP has previously been shown to elicit antibodies predominantly targeting the GC and MLD [31,32]. These antibodies are generally non-neutralizing as both domains get cleaved off during EBOV entry into cells before receptor binding and fusion occurs [33]. However, with the increase in antibody-based therapeutics development a few GC domain-binding antibodies have been described to potently neutralize EBOV infection and to be polyfunctional [34,35], which is in general associated with increased vaccine efficacy [36]. In addition, the monoclonal antibody cocktail ZMapp contains the GC-binding antibody 13c6 [37]. Recently, inhibition of cathepsin cleavage by these GC-binding antibodies was proposed as a mechanism [38]. Therefore, while a deletion of the GC and MLD might not interfere with the GP’s ability to drive VSV replication, it could result in reduced protective immunity against EBOV infection. We compared the VSV vectors expressing EBOV GP [8], the EBOV GPΔMLD [17], and the EBOV GPΔGCΔMLD, finding no significant differences in replication in cell culture (Fig. 1C), which highlights that both domains, GC and MLD, are indeed dispensable for *in vitro* replication of the VSV vectors.

Ultimately, we wanted to know if a single dose vaccination with the VSV-EBOVΔGCΔMLD or VSV-EBOVΔMLD resulted in protection in mice as has been described for VSV-EBOV [39]. We observed that nearly all (5/6) mice vaccinated with VSV-EBOVΔGCΔMLD succumbed to EBOV infection, whereas all mice vaccinated with VSV-EBOVΔMLD survived without signs of disease (Fig. 2C). When comparing virus loads in target tissues like spleen and liver, we found that the titers in the VSV-EBOVΔGCΔMLD group were similar to control-vaccinated animals and significantly higher than in mice vaccinated with the other two vaccines (Fig. 2D). Comparing virus titers on day 4 post-challenge between the VSV-EBOV- and VSV-EBOVΔMLD-vaccinated mice, we only observed a significant difference in viremia (Fig. 2D). This difference might explain the slightly higher weight gain after challenge in the VSV-EBOV compared to VSV-EBOVΔMLD-vaccinated mice (Fig. 2B).

Analysis of humoral immune responses after vaccination and challenge supported the observed survival differences. While it was not a significant difference, the EBOV GP-specific IgG on the day of challenge was lower in the VSV-EBOVΔGCΔMLD compared to the VSV-EBOV- and VSV-EBOVΔMLD-vaccinated groups (Fig. 3A). On day 4 after challenge the VSV-EBOVΔGCΔMLD group had IgG levels comparable to controls indicative of antibody-consumption by virus replication. Interestingly, we observed a small decline in IgG levels also in the VSV-EBOVΔMLD-vaccinated mice on day 4 possibly indicating antibody consumption by virus replication as evidenced by viremia (Fig. 2D). The challenge virus replication in the VSV-EBOVΔMLD-vaccinated mice resulted in a boost effect of the humoral immune response resulting in an increase in particularly the neutralizing titers form ~1:160 on day 0 (Fig. 3B) to 1:2,560 on day 42 (Fig. 3C). Similarly, the neutralizing titer in the single surviving mouse in the VSV-EBOVΔGCΔMLD group increased. In contrast, the neutralizing activity in the serum from VSV-EBOV-vaccinated and surviving mice remained the same or slightly decreased by day 42 (Fig. 3B, C) because the vaccine-induced immune responses controlled challenge virus replication as shown by no viremia (Fig. 2D) and, therefore, no boost effect was observed.

In summary, our data support and expand existing data sets about the importance of antibodies directed to the EBOV GP GC for protection from EVD [31,32,38]. Our findings also demonstrate that both the GC and MLD are dispensable for VSV-EBOV-based vaccine vector replication *in vitro*. If the EBOV GP is only used to target important immune cells (“vehicle”), a VSV vector expressing the EBOV GPΔGCΔMLD might be of advantage as the immune response will likely be biased, as intended, towards the second foreign immunogen. However, when developing a truly bivalent vaccine with the aim to similarly protect against EBOV and a second pathogen, a VSV vector expressing EBOV GPΔMLD or full-length EBOV GP would be advantageous to not lose protective immune responses against EBOV. Future studies will focus on directly comparing the protective efficacy of VSV-EBOV- and VSV-EBOVΔGCΔMLD-based vaccines in order to optimize second generation VSV-based vaccine vectors.

## Acknowledgements

We thank the animal care takers of the Rocky Mountain Veterinary Branch (NIAID) for their support of the mouse study and Josephin Greßler (FLI) for technical assistance generating the tetracistronic minigenome plasmid.

## Author contributions

H.F. and A.M. conceived the idea and secured funding. A.M. designed the studies. B.B. generated the VSV-EBOVΔGCΔMLD vaccine and performed ELISAs. B.B., W.F. and A.M. performed the mouse study. T.H. established the tetracistronic minigenome plasmid. W.F. performed the neutralization assays. A.M. wrote the manuscript with contributions from all authors. All authors approved the manuscript.

